# *MedGraphNet* : Leveraging Multi-Relational Graph Neural Networks and Text Knowledge for Biomedical Predictions

**DOI:** 10.1101/2024.09.24.614782

**Authors:** Oladimeji Macaulay, Michael Servilla, David Arredondo, Kushal Virupakshappa, Yue Hu, Luis Tafoya, Yanfu Zhang, Avinash Sahu

## Abstract

Genetic, molecular, and environmental factors influence diseases through complex interactions with genes, phenotypes, and drugs. Current methods often fail to integrate diverse multi-relational biological data meaningfully, limiting the discovery of novel risk genes and drugs. To address this, we present *MedGraphNet*, a multi-relational Graph Neural Network (GNN) model designed to infer relationships among drugs, genes, diseases, and phenotypes. *MedGraphNet* initializes nodes using informative embeddings from existing text knowledge, allowing for robust integration of various data types and improved generalizability. Our results demonstrate that *MedGraphNet* matches and often outperforms traditional single-relation approaches, particularly in scenarios with isolated or sparsely connected nodes. The model shows generalizability to external datasets, achieving high accuracy in identifying disease-gene associations and drug-phenotype relationships. Notably, *MedGraphNet* accurately inferred drug side effects without direct training on such data. Using Alzheimer’s disease as a case study, *MedGraphNet* successfully identified relevant phenotypes, genes, and drugs, corroborated by existing literature. These findings demonstrate the potential of integrating multi-relational data with text knowledge to enhance biomedical predictions and drug repurposing for diseases.*MedGraphNet* code is available at https://github.com/vinash85/MedGraphNet

## 1. Introduction

Diseases such as cancer are multifaceted, influenced by genetic, molecular, and environmental factors, as well as their interactions. Most diseases have a genetic component [13], where genotype interacts with environmental factors like lifestyle, influencing the manifestation of an individual’s phenotype or disease. The phenotype remains relatively stable throughout life, unless altered by environmental factors or disease interactions with genotype [43]. Additionally, gene-gene interactions, or epistasis, can lead to drug or treatment failures [35]. Thus, complex interactions exist between diseases, genes, phenotypes, and drugs.

Diseases, drugs, phenotypes, and genes have each been studied extensively. Research is usually focused on dissecting these complexities into individual relationships: disease-drug, disease-gene, disease-phenotype, phenotype-gene, phenotype-drug (drug side effects), gene-drug, and gene-gene interactions, resulting in extensive databases with millions of interactions [42, 27]. Integrating these disparate components may reveal new drug targets for specific diseases [33]. Experimental discovery of gene associations is typically carried out through genome-wide association studies (GWAS), which often fail to detect associations for rare or uncommon diseases due to unique or less common genetic mutations that are not adequately captured [38, 5]. GWAS have reported over 50,000 gene-disease associations [38, 5], but require large cohorts for statistical power to detect small associations [38, 5]. Complex diseases like schizophrenia, obesity, asthma, and hypertension involve gene-environment interactions, complicating the identification of genetic bases and effective drugs. Polygenic diseases also present difficulties in detecting genetic associations [14].

Despite progress, current methods fail to integrate diverse biological data meaningfully, obstructing the discovery of novel risk genes and drugs. Methods using relational data often rely on a relation using two types of nodes, limiting their effectiveness. Such models, called Single Relation Graph (SRG) models, are insufficient in various scenarios, such as rare diseases, where conducting genome-wide association studies is challenging due to small sample sizes and the absence of single risk factors for imputation. Graph Neural Networks (GNNs) excel at predicting new interactions within a network of known connections. However, SRGs, which typically initialize nodes randomly [3], depend solely on connections for inference and struggle to learn representations for isolated or sparsely connected nodes.

We therefore present *MedGraphNet*, a GNN-based approach to infer relationships between biological entities: drug, gene, disease, and phenotype leveraging multi-relational data. Nodes are initialized with highly informative embeddings created from GeneLLM [14] representations of text summaries, allowing generality across data types and facilitating a more transferable model. By incorporating multiple modalities or node types, the graph infers connections for isolated nodes through indirect paths not possible in SRG models. This enhanced connectivity allows the GNN to predict interactions of any type (disease-drug, disease-gene, etc.) by learning from existing knowledge and multi-relational interactions. This is particularly useful for predicting unknown risk genes and drug-disease associations in rare diseases. Therefore, we tested the hypothesis that *MedGraphNet* will outperform traditional single-relation graph methods in predicting associations between biomedical entities, and showed that it can accurately infer risk genes for diseases, suggest new drugs for untreated diseases, infer potential side effects of drugs, and provide interpretable insights into biological mechanisms. This approach addresses the critical gap of integrated, multi-relational analysis, with the potential to advance actionable biomedical predictions.

Related works can be found in Appendix **A**.

## 2. Methods

### 2.1. Data

For training *MedGraphNet*, we compiled datasets from several databases, which we collectively refer to as MedGraphDB (Table 1). In addition to databases contained in MEd-GraphDB, we also collected additional databases for independent evaluation of *MedGraphNet* (Table 1).

**Table 1:** Source and Graph statistics of MedGraphDB and other evaluation datasets.

| Data | Association | Source | Num of Nodes Used | Num of Edges |
| --- | --- | --- | --- | --- |
| **MedGraphDB** | Disease-gene | MGI [20] | 1839 Diseases & 2897 Genes | 8232 |
|  | Gene-Drug | DGIdb [7] | 2101 genes & 2814 Drugs | 21,080 |
|  | Disease-Drug | BIOSNAP [21] | 1985 Diseases & 1070 Drugs | 142,346 |
|  | Phenotype-Gene | HPO [18] | 9497 Phenotypes & 4540 Genes | 827,172 |
|  | Disease-Phenotype | HPO [18] | 4809 Diseases & 6599 Phenotypes | 96,693 |
|  | Gene-Gene | PPI [37] | 14060 Genes | 344,638 |
| **Other Evaluation Datasets** | Disease-gene | DisGeNET [28] | 986 Diseases & 5405 Genes | 16,849 |
|  | Disease-gene | PsyGeNET [8] | 16 Diseases & 1131 Genes | 3,910 |
|  | Phenotype-Drug | SIDER 4.1 [19] | 713 Phenotypes & 800 Drugs | 34701 |

#### 2.1.1. MedGraphDB compilation

MedGraphDB includes disease-drug, phenotype-gene, disease-gene, disease-phenotype, gene-drug, gene-gene, and drug-phenotype associations, summarized in Table 1. For additional details, refer to Appendix B. Note that, drug side-effect relational database was purposely not included in MedGraphDB.

In MedGraphDB, we also compiled summaries of diseases, genes, drugs, and phenotypes from the Human Disease Ontology [32], UNIPROT [4], PubChem [16], and Wikipedia databases, respectively as described in the Appendix B.

### 2.2. MedGrapNet

MedGrapNet is a heterogeneous graph consisting of four types of nodes: (diseases, genes, drugs, and phenotypes), and edges between them depicting their known relationships. The node indexing and edge construction steps can be found in Appendix C and D. *MedGraphNet* consists of 32,940 nodes and 1,440,161 edges. The number of nodes and edges of each types is summarized in Tables 1 and Appendix E.

#### 2.2.1. Initialization

Nodes of *MedGraphNet* were initialized using summaries from MedGraphDB. The summaries are converted into embeddings using a GeneLLM [14], which outputs embedding of fixed size from text summaries as input. GeneLLM [14] is an interpretable transformer-based model trained on both unstructured textual information and structured Gene Ontology (GO) relationships using contrastive learning. GeneLLM embeddings were obtained for diseases, drugs, phenotypes, and genes using their text summaries from MedGraphDB.

#### 2.2.2. Handling Heterogeneous Graph Neural Network using GCN

All nodes are initialized using GeneLLM’s 768-dimensional vectors, but embeddings for diseases, drugs, phenotypes, and genes reside in type-specific spaces, forming a heterogeneous graph. We compared various heterogeneous graph neural networks, such as HAN [41], HetGNN [45], and R-GCN [31], and selected the approach by Qingsong *et al.* [25], which extends GCNs for heterogeneous graphs using type-specific fully connected layers. These layers process and adjust feature representations for each node type, co-embedding them in a common latent space while preserving type-specific information.

To unify the embedding representations, we use type-specific fully connected layers trained end-to-end with GCN parameters. The *MedGraphNet* architecture consists of multiple layers of GCN aggregate and transform features from adjacent nodes as 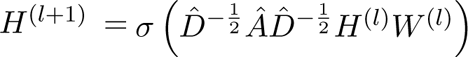 where *H*^(*l*)^ is node embeddings at layer *l*, 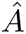 is the adjacency matrix, 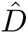 is degree matrix, *W* ^(*l*)^ is the trainable weight matrix, and *σ* is the ReLU activation (details in Appendix F.0.1).

##### Training

The model was optimized for binary cross entropy loss using the Adam optimizer with an initial learning rate of 0.01 and weight decay, and early stopping was used to prevent overfitting (Appendix F). The GNN is first initialized with all nodes. The set of known edges are referred to as positive samples, and labeled as 1. In each set of samples (test/train/validation), an equal number of negative samples were created by randomly selecting an edge between two non-connected nodes and labeled 0. These sets of positive and negative edges are then used to train the model for the task of predicting whether or not an edge exists between two nodes. First we removed all phenotype-drug edges (drug side effect) from the dataset, in order to test the model’s ability to predict edges with completely unknown associations. Then we trained a model, termed Random Split, by randomly splitting the list of all remaining edges into training (80%), validation (10%), and test sets (10%).

In addition to this model, we also created hard test sets, in which all of a given node’s edges for a particular association are only present in the test set, i.e., an *isolated node*. For example, a given disease could be isolated from all gene nodes in training, forcing the model to learn these edges based only on its connections to phenotype and drug nodes. To create this dataset, we first split the nodes into 80/10/10 groups. Then to create an isolated test set for gene-drug associations, all positive samples for that gene are placed in the test set. This means that the distribution of edges may not be 80/10/10 as in the randomly sampled training set. *Leaf node* test sets were also created this way by placing one edge for a given node into the training set. For each association type there can be two isolated or two leaf node sets created (i.e., isolated gene or isolated drug in gene-drug association).

##### Cluster Analysis

First tSNE clusters were created for the refined embeddings from the trained Random Split model. Over Representation Analysis (ORA) is performed using the WebGestaltR package [23] to identify enriched pathways for each of the gene clusters. To further explore the relationships between clusters, we then performed a hypergeometric test to identify significant associations between all clusters of the nodes based on the model’s predictions. The hypergeometric test assesses whether the association between the nodes in two clusters is greater than expected by chance, and all associations between clusters with p-value less than 0.05 after adjusting for multiple hypothesis are considered significant.

### 2.3. Baselines

To our knowledge, no existing methods predict all six relationship types (Disease-Phenotype, Drug-Disease, Gene-Disease, Gene-Drug, Phenotype-Gene, and Phenotype-Drug) that *MedGraphNet* addresses. To ensure uniform comparison across relationship types, we bench-marked *MedGraphNet* against two baselines: *MedGraphNet* Single Relation Graph (*MedGraphNet-SRG*) and Multimodal Fusion.

#### Multimodal Fusion

For each relationship type, we trained a separate fusion model using GeneLLM embeddings as inputs. For instance, in predicting Drug-Gene relations, we started with GeneLLM embeddings of drugs and genes. We fused these embeddings and applied a series of fully connected layers, culminating in a binary output that classifies whether a positive link exists between the drug-gene pair in the training data. The model was trained to minimize the cross-entropy loss between predicted and actual links, using the same data as *MedGraphNet* for a fair comparison. This approach was similarly applied to other relationship types.

#### MedGraphNet Single Relation Graph (MedGraphNet-SRG)

For each relationship type, we trained a GNN using a graph comprising only that type’s edges. For example, for Drug-Gene relations, we applied the GNN to a graph with nodes representing drugs and genes, and edges representing their relations. The training and testing procedures for *MedGraphNet-SRG* were similar to those used for *MedGraphNet*.

## 3. Results

### 3.1. Baselines

To evaluate the performance of *MedGraphNet*, we compared its performance to both single-relation graph (SRG) and multimodal fusion methods using different test sets. The test sets are grouped based on whether a node’s edge is isolated completely from the training data, or it has only one edge in the training set (leaf node). The AUC of the models on the different test cases is summarized in Tables 1 - 6.

#### Gene-Drug Association

For Gene-Drug association tasks, *MedGraphNet* demonstrated superior performance for test cases with 10% of the nodes isolated nodes (Methods), whereas *MedGraphNet-SRG* exhibited higher performance for random test cases. In scenarios with isolated nodes, the absence of edges connecting isolated genes to drugs in the training set prevented *MedGraphNet-SRG* from propagating information to these isolated nodes. In contrast, *MedGraphNet* overcame the lack of edges between isolated nodes by propagating information between edges of isolated nodes and genes or phenotypes. Consequently, *MedGraphNet* outperformed *MedGraphNet-SRG* for isolated nodes (Table 2).

**Table 2:** Comparison for Gene-Drug Association. Highest performance in bold.

| Test Case | *MedGraphNet* | SRG | Multimodal Fusion |
| --- | --- | --- | --- |
| Gene Isolated Node | **0.95** | 0.31 | 0.85 |
| Drug Isolated Node | **0.92** | 0.81 | 0.85 |
| Random Split | 0.91 | **0.97** | 0.94 |

To investigate if this limitation of SRG is also evident when there are few edges connected to nodes, we studied gene or drug nodes with a single edge in the training set (referred as leaf node). Performance for leaf nodes was also lower for SRG than for *MedGraphNet* (Table G.1), suggesting that connectivity significantly impacts model performance. Interestingly, in the Random Split test case, SRG performed the best.

#### Disease-Gene, Disease-Phenotype, Disease-Drug, Phenotype-Gene Associations

All tasks showed similar performance trends as observed for Disease-drug associations, as illustrated in Table 2. In all cases, GNN performance depended on connectivity, with *MedGraphNet* outperforming SRG in all instances (Appendix G.2, G.3, G.4, and G.5).

*MedGraphNet-SRG* baseline method for Disease-Gene Association prediction is exactly equivalent to the method recently proposed by Rifaioglu *et al*. ^1^. Consistent with our results reported in Appendix G.2, their method also reported high accuracy for Disease-Gene prediction. However, these SRG methods struggle in challenging test cases with isolated nodes, which is particularly important for predicting rare diseases without any known gene associations. Additionally, as shown in Section 3.2, the SRG-based method performs poorly when trained in one cohort and applied to another cohort.

### 3.2. Model Generalizability

To verify the generality of *MedGraphNet*, we implemented three assessment methods:

1. Drug-Phenotype Relationships: MedGraphDB excluded known Drug-Phenotype relationships, meaning *MedGraphNet* was not trained on them. We investigated whether *MedGraphNet* could infer these relationships independently. As shown in (Figure 2a), *MedGraphNet* accurately inferred Drug-Phenotype relationships. Remarkably, it even outperformed a GNN model directly trained on Drug-Phenotype relationships, achieving superior prediction accuracy in 20% of the test data. This underscores the advantage of a multi-relational graph and its ability to generalize to new relationship types.
2. Disease-Gene Associations: We predicted disease-gene associations using DisGeNET [28] and PsyGeNET datasets [8], which are external to MedGraphDB and not included in training. Unlike the MGI diseases dataset [20] with 8,231 edges between diseases and genes, DisGeNET [28] and PsyGeNET [8] contain 16,850 and 3,911 edges, respectively. *MedGraphNet* consistently achieved an AUC of 0.87 and 0.88 for both DisGeNET [28] and PsyGeNET [8], respectively, compared to SRG and Multimodal Fusion as shown in (Figure 2b), demonstrating robust generalizability of MedGraphDB to unseen datasets.
3. Performance on Rare Diseases: A major challenge for many rare diseases is that no disease-gene associations are known. To evaluate *MedGraphNet* ’s performance on such diseases, we simulated a rare disease scenario by randomly selecting diseases from MedGraphDB and removing all edges between these diseases and any genes from the training set. We then retrained *MedGraphNet* with this new graph. Remarkably, the model maintained an AUC of 0.94 for these selected diseases, similar to its performance on diseases with known associations (Figure 2c). This suggests that for diseases without known risk genes, *MedGraphNet* can leverage edges connected to diseases with known phenotypes and drugs to infer associations.

**Figure 1:**
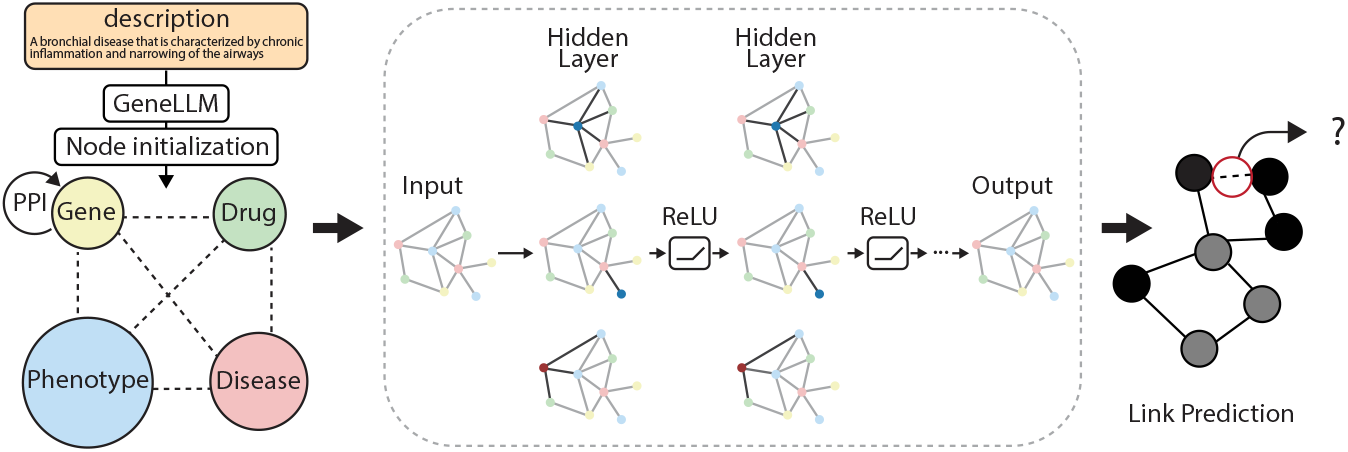
An overview of the *MedGraphNet* link prediction model architecture showing node initialization using GeneLLM embeddings and the interactions between genes, drugs, phenotypes, and diseases.

**Figure 2:**
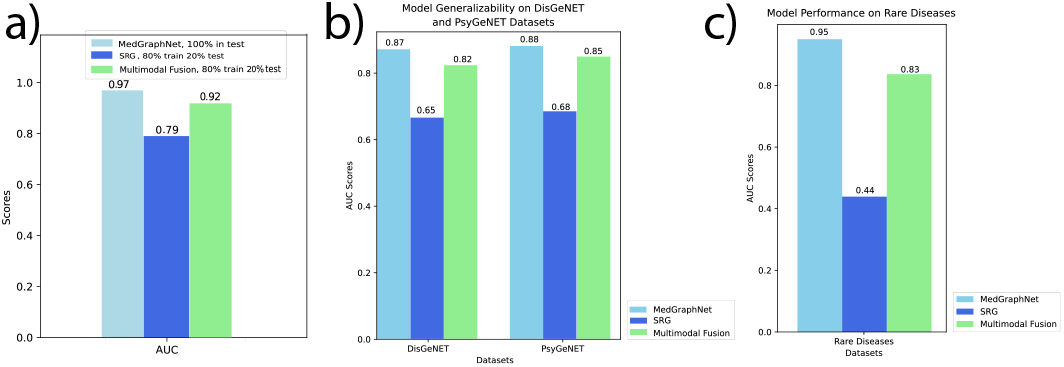
Model Generalizability of *MedGraphNet* : (a) *MedGraphNet* ’s performance in inferring Drug-Phenotype relationships, (b) Comparison of AUC scores on DisGeNET and PsyGeNET datasets, (c) Comparison of AUC scores for rare diseases, demonstrating *MedGraphNet* ’s ability to generalize to unseen datasets.

### 3.3. Applications

#### 3.3.1. Alzheimer’s disease as Case Study

Using *MedGraphNet*, we asked if given a disease, can we correctly drugs associated with the disease.

Using Alzheimer’s disease as a case study, the model was able to infer meaningful associations. Predicted phenotype associations included mental deterioration, cognitive fatigue, and emotional lability, supported by studies from Knopman et al. [17], Angioni et al. [1], and Silva et al. [34]. Table 3 shows the top 3 predictions for each of the associations, and supporting literature.

**Table 3:** Top Predictions for Alzheimer’s Disease Aspredict phenotypes, genes, and sociations and Supporting Literature.

| Alzheimer’s | Associate | Literature |
| --- | --- | --- |
| Phenotype | Mental deterioration | Knopman *et al* [17] |
|  | Cognitive fatigue | Angioni *et al* [1] |
|  | Emotional lability | Silva *et al* [34] |
| Genes | PICALM | Harold *et al* [9] |
|  | TREM2 | Jonsson *et al* [15] |
|  | EPHA1 | Hollingsworth *et al* [11] |
| Drugs | Namzaric | Tariot *et al* [40] |
|  | Donepezil | Salloway *et al* [30] |
|  | Solanezumab | Honig *et al* [12] |

#### 3.3.2. Cluster analysis

Next, we inferred clusters for each biological entity using the node embeddings (Figures 3b, G.1, G.2, G.3) and examined the relationships between these clusters. We hypothesized that *MedGraphNet* could help identify the underlying mechanisms that make certain drug clusters effective treatments for specific disease clusters. To test this hypothesis, we created a cluster graph (Figure 3b), connecting two clusters if the number of known inter-cluster edges was significantly higher than expected by random chance (Section 2.2.2).

**Figure 3:**
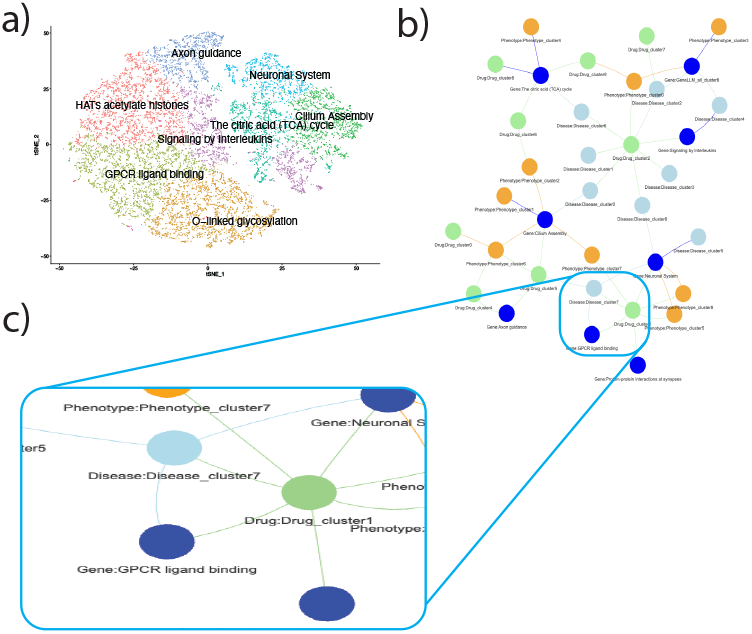
*MedGraphNet* node cluster annotation. a) t-SNE Plot of Gene Embeddings Clustering with Enriched Pathway Names. Each cluster is labeled by the most significantly enriched pathway. b) Network graph showing significant association between clusters of gene, disease, drug and phenotype

The cluster graph revealed that the molecular basis of drug clusters could be linked to disease clusters through their association with common gene clusters. For example, the GPCR ligand binding gene cluster forms a clique with Disease Cluster 7 and Drug Cluster 1 (Figure 3 c). Part of the drugs in drug cluster 1 is *Acamprosate*. One of the drugs in Drug Cluster 1 is Acamprosate, which inhibits the trans-ACPD binding resulting in diminished neurotoxicity, indicating an interaction with metabotropic glutamate receptors (a type of GPCR) (Harris et al., 2002) [10]. Acamprosate is also linked to Alcohol Use Disorder, a disease found in Disease Cluster 7, and has been reported as an effective treatment for this condition (Maisel *et al* in 2013)[26]. Another example is Risperidone that is used to treat schizophrenia (Cluster 7). Risperidone can cause an irreversible interaction with h5-HT7 receptors, a family of GPCRs [36]. These examples demonstrate how the node embeddings can reveal meaningful connections between drugs, diseases, and genes.

### 3.4. Ablation Study

The comparison between the fusion model and *MedGraphNet* demonstrated that incorporating relational information through a GNN is crucial for accurate prediction. To assess the necessity of text summaries, we compared *MedGraphNet* with its variant that initializes nodes randomly (*Med-GraphNet* -Random). The random initialization exhibited lower performance (Appendix G.6), underscoring the importance of using text summaries for node initialization to achieve high predictive accuracy.

## 4. Conclusion

*MedGraphNet*, a multi-relational GNN-based model, demonstrates promising improvements over traditional single-relation graph methods and multimodal fusion approaches in predicting associations among biomedical entities: genes, drugs, diseases, and phenotypes. It addresses the challenge of isolated or sparsely connected nodes by utilizing existing text knowledge through an LLM and leveraging multi-relational data. Our results indicate that *MedGraphNet* trained on one cohort can generalize to new, unseen cohorts and identify risk genes for rare diseases. Remarkably, despite not being trained on drug side-effect data, *MedGraphNet* could predict side effects, in some cases outperforming models directly trained on this data. Using Alzheimer’s disease as a case study, *MedGraphNet* identified risk genes and drug associations not present in the training data but supported by existing literature, highlighting its potential to uncover insights for rare diseases. Additionally, *MedGraphNet* proved useful for identifying and annotating clusters of drugs and diseases based on shared genetic mechanisms. These findings suggest that *MedGraphNet* ’s multi-relational framework and use of informative embeddings can enhance biomedical predictions, providing valuable insights into disease mechanisms and potential drug repurposing strategies. However, experimental validation is necessary to confirm these predictions and advance its clinical utility.

## Appendix A. Related Research

### Graph Neural Networks

Multimodal knowledge graphs are being used to gain better insights into complex medical diseases [44]. Graph neural networks have been applied to computational biology projects since 2019 [46]. By examining hidden relationships between distinct entities such as drugs, diseases, and genes, and analyzing the topology and structure of these entities, personalized medicine can be improved, including for patients with complex diseases [44]. Several studies have leveraged heterogeneous graph machine learning tools to health-related contexts.

Long et al. [24] utilized a heterogeneous graph attention network to predict SARS-CoV-2 drug-virus associations, integrating diverse biomedical data sources into two heterogeneous graphs for drug-virus/microbe associations and drug-target interactions, achieving state-of-the-art performance. Asada et al. [2] combined heterogeneous pharmaceutical data to predict drug-drug interactions, demonstrating superior performance in DDI extraction and drug-protein interactions, highlighting the advantage of applying GNNs on multirelational data over unirelational data. Tanvir et al. [39] introduced a triplet attention mechanism within a heterogeneous graph to model drug-target-disease interactions.

### Drug-target predictions

Several machine learning approaches have been used to study and predict drug-target interactions (DTIs) using features derived from molecular structures, biological activity, and other relevant data. Early studies includes SVMs to predict protein-chemical interactions based on amino acid sequences, chemical structures, and mass spectrometry data [29]. Additionally, random forest techniques were used to combine non-structural descriptors of drugs or chemicals and their targets in a proteochemometric modeling approach [29]. Recently, deep learning methods have also been deployed for DTI prediction[29]. GNN has been recently applied for DTI prediction and shown to outperform other approaches [47, 22].

### Drug side-effect prediction

Various machine learning methods have been applied to the prediction of drug side effects. Semi-supervised methods such as clustering, traditional supervised machine learning approaches, and deep learning techniques have all been employed [6]. However, class imbalance remains a significant challenge in multi-task drug side-effect prediction, necessitating new methods.

## Appendix B. Datasets

### Disease-Gene Association Dataset

The dataset was obtained from the DisGeNET database [28]. DisGeNET [28] is a comprehensive discovery platform containing one of the largest publicly available collections of genes and variants associated with human diseases. It integrates data from expert-curated repositories, genome-wide association studies (GWAS) catalogues, animal models, and the extensive corpus of scientific literature. We downloaded the Gene-Disease Associations (GDAs) dataset, which contains 1,134,942 GDAs, linking 21,671 genes with 30,170 diseases, disorders, traits, and phenotypes. Data cleaning involved removing duplicate entries and reperesenting genes and diseases with their IDs (https://www.disgenet.org/).

### Phenotype-Gene and Phenotype-Disease Association Datasets

The dataset was sourced from the Human Phenotype Ontology (HPO) [18]. This dataset includes phenotype-gene pairs, representing associations between observed phenotypes and genetic factors. It comprises 3,000 phenotype-gene pairs. It also consists of 4,500 disease-phenotype pairs, capturing the association of phenotypes to diseases.

### Disease-Drug Association Dataset

The dataset was obtained from the Stanford Network Analysis Project (SNAP), specifically from the BIOSNAP Datasets [21]. It is a disease-drug association network that contains information on drug-disease relationships. Nodes represent diseases and drugs (also including certain chemicals that are not human drugs). It contains 1,535 diseases, 1,662 chemicals and 466,656 associations. The chemicals that are not human drugs were filtered out.

### Gene-Drug Association Dataset

The dataset was obtained from the Drug-Gene Interaction Database (DGIdb) [7]. DGIdb integrates drug-gene interactions mined from various sources such as DrugBank, PharmGKB, ChEMBL, and Drug Target Commons. It includes over 10,000 genes and 15,000 drugs involved in over 50,000 drug-gene interactions. Data were cleaned by removing duplicate entries and using drug and gene IDs.

### B.1. Node Summaries

We compiled summaries of diseases, genes, drugs, and phenotypes from several sources, as described below.

#### Human Disease Ontology

The Human Disease Ontology (DO) [32] is a standardized ontology for human diseases. It provides descriptions of human disease terms. We utilized the DO to extract and compile disease summaries.

#### UNIPROT

UNIPROT [4] is a comprehensive resource for protein sequence and functional information. We used gene summary data from UNIPROT to generate the gene node embeddings

#### PubChem

PubChem [16] is a public repository for information on the biological activities of small molecules. We used PubChem to gather data summaries of various drugs, including their chemical properties.

#### Wikipedia

Wikipedia is a widely-used, crowd-sourced encyclopedia. We extracted phenotypic information from Wikipedia to complement the data from other sources.

## Appendix C. Node Identification and Indexing

To construct the graph, we first identified and indexed the nodes from all the datasets. The nodes in our graph represent diseases, drugs, phenotypes, and genes. Nodes representing diseases were identified from the disease-related datasets, including disease-drug, disease-gene, and disease-phenotype datasets. Drugs were identified from the disease-drug and gene-drug datasets. Phenotypes were sourced from the phenotype-related datasets, and the genes were also identified in a similar manner. Each type of node was then assigned a unique index to ensure proper integration into the graph structure. Diseases were indexed starting from 0. Drugs were indexed starting from the end of the disease indices. Phenotypes were indexed starting from the end of the drug indices, and genes were indexed starting from the end of the phenotype indices.

## Appendix D. Edge Construction

Edges in the graph represent various types of associations between the nodes. We constructed edges based on the pairs present in each dataset. For example, in the disease-drug dataset, an edge was created between the corresponding disease node and drug node for each pair in the dataset. The edge construction ensures that all relevant associations are represented in the graph, for effective analysis of the relationships between the different entities (disease, drug, gene, and phenotype).

## Appendix E. Graph Statistics

The constructed graph comprises a total of 32,940 nodes and 1,440,161 edges. Specifically, the graph includes 5,968 disease nodes, 3,212 drug nodes, 9,597 phenotype nodes, and 14,163 gene nodes. The edges represent various associations: 142,346 disease-drug, 827,172 phenotype-gene, 8,232 disease-gene, 96,693 disease-phenotype, and 21,080 gene-drug interactions, as shown in (Table 1).

**Table E.1:**
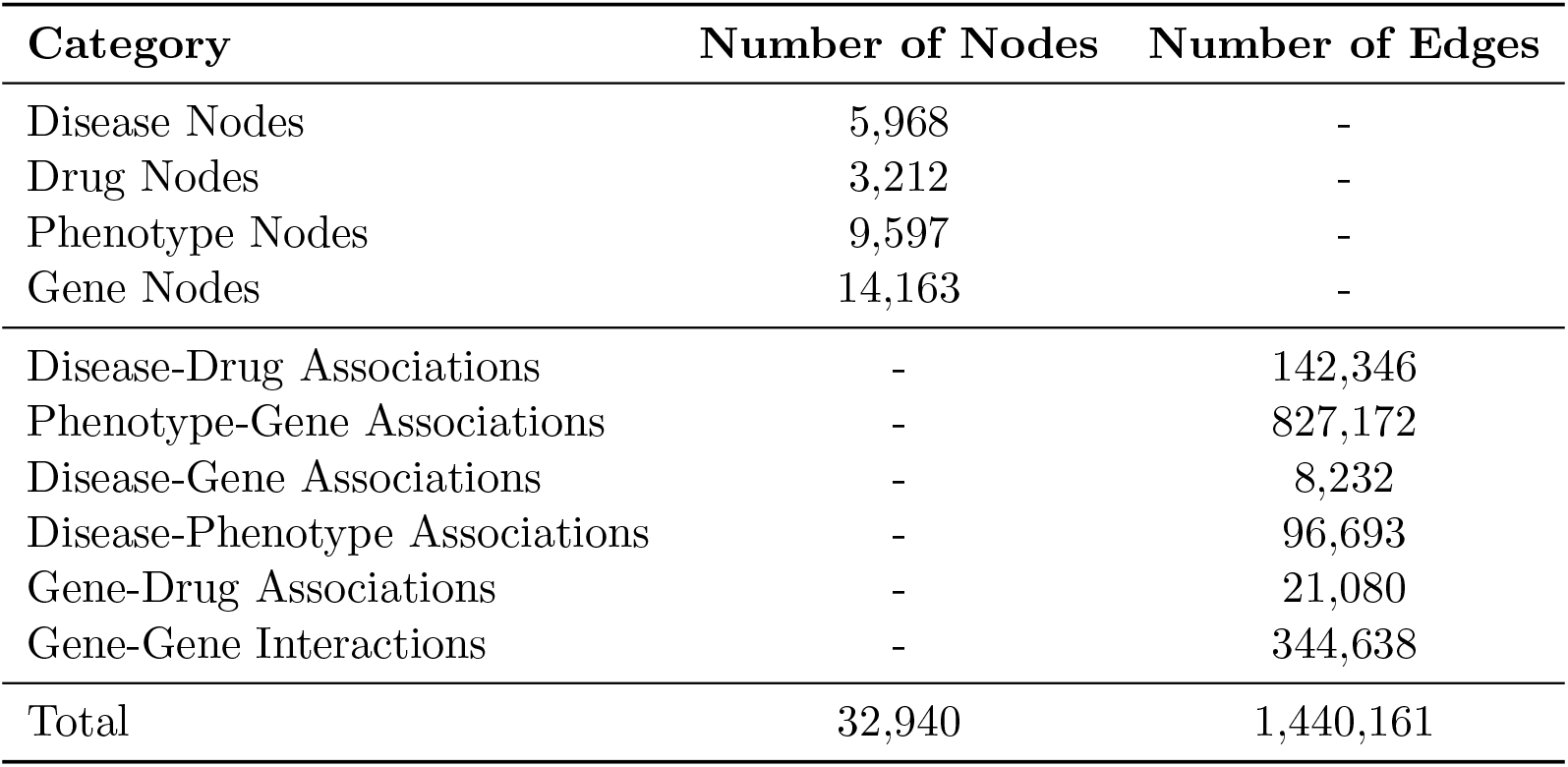
Graph Statistics.

## Appendix F. Training

- **Input Layer:** The graph data consisting of node embeddings, edges, optional edge weights, and labels corresponding to each node are taken as input.
- **Graph Convolution Layer:** Multiple layers of graph convolution operations are applied to aggregate and transform features from adjacent nodes. The convolution

operation is defined as:

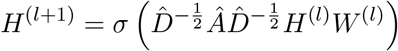

where *H*^(*l*)^ represents the node embeddings at layer *l*, 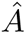 is the normalized adjacency matrix, 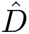 is the diagonal degree matrix, *W* ^(*l*)^ is the layer-specific trainable weight matrix, and *σ* is the ReLU activation function.

- Each convolution layer is followed by an activation function to introduce non-linearity to the network and a dropout regularization function to randomly drop a portion of the node features during training to reduce overfitting.
- **Output Layer:** The final features are processed through a linear layer or convolution layer to produce the final node embeddings that incorporate information propagated through the graph.

By adjusting parameters such as the number of layers and the dimensionality of the hidden layers, the model can be adapted to different scales of graphs and complexities of tasks.

### F.0.1. Loss Function and Optimization

To train the GNN model, we used a loss function suitable for binary classification, given that the task involves predicting the existence of an edge (association) between nodes.

#### Binary Cross-Entropy Loss

We employed the Binary Cross-Entropy (BCE) loss function, which is defined as:

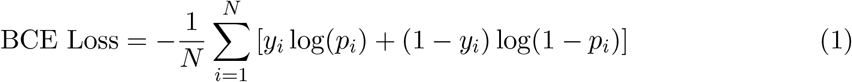

where *N* is the number of samples, *y_i_*is the true label (1 for positive samples, 0 for negative samples), and *p_i_* is the predicted probability of the edge’s existence.

#### Optimization

The optimization of the GNN model was carried out using the Adam optimizer, which is well-suited for training deep learning models due to its adaptive learning rate capabilities. The key parameters of the optimizer included:

- **Learning Rate**: The initial learning rate was set to 0.01.
- **Weight Decay**: A small weight decay value was used to prevent overfitting.

During each training epoch, the model’s parameters were updated by minimizing the BCE loss, and the training process continued until the validation loss stopped improving, indicating convergence.

## Appendix G. Results

**Table G.1:** Performance Metrics of models trained with Leaf Node Association. Bold values indicate the highest performance among the methods.

| Association | Test Case | *MedGraphNet* | SRG |
| --- | --- | --- | --- |
| Gene-Drug | Gene Leaf Node | **0.95** | 0.79 |
|  | Drug Leaf Node | **0.93** | 0.85 |
| Disease-Gene | Disease Leaf Node | **0.94** | 0.77 |
|  | Gene Leaf Node | **0.92** | 0.74 |
| Disease-Drug | Disease Leaf Node | **0.85** | 0.75 |
|  | Drug Leaf Node | **0.96** | 0.93 |
| Phenotype-Gene | Phenotype Leaf Node | **0.87** | 0.67 |
|  | Gene Leaf Node | **0.88** | 0.67 |
| Disease-Phenotype | Disease Leaf Node | **0.90** | 0.80 |
|  | Phenotype Leaf Node | **0.93** | 0.90 |

**Table G.2:** Performance Metrics Comparison for Disease-Gene Association. Bold values indicate the highest performance among the three methods.

| Test Case | *MedGraphNet* | SRG | Multimodal Fusion |
| --- | --- | --- | --- |
| Disease Isolated Node | **0.95** | 0.44 | 0.83 |
| Gene Isolated Node | **0.95** | 0.73 | 0.88 |
| Random Split | 0.94 | **0.98** | 0.92 |

**Table G.3:** Performance Metrics Comparison for Disease-Drug Association. Bold values indicate the highest performance among the three methods.

| Test Case | *MedGraphNet* | SRG | Multimodal Fusion |
| --- | --- | --- | --- |
| Disease Isolated Node | **0.89** | 0.63 | 0.66 |
| Drug Isolated Node | **0.98** | 0.92 | 0.61 |
| Random Split | **0.94** | 0.80 | 0.90 |

**Table G.4:**
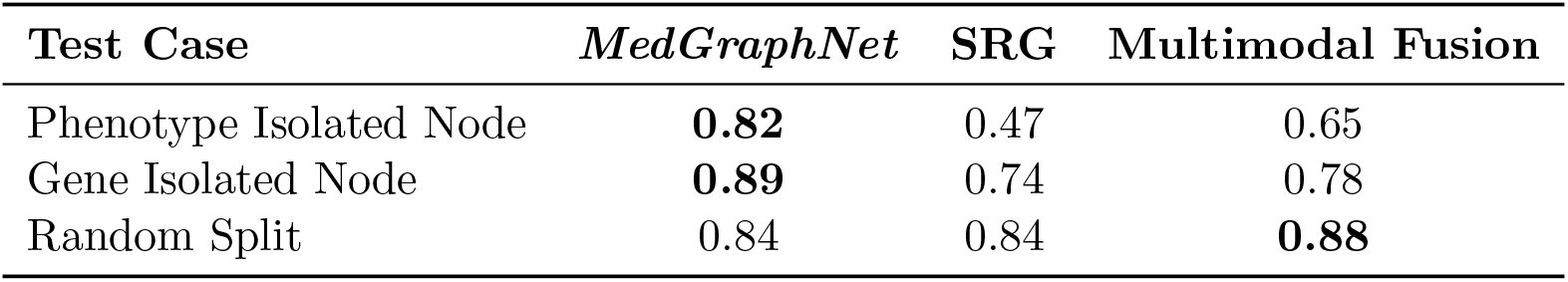
Performance Metrics Comparison for Phenotype-Gene Association. Bold values indicate the highest performance among the three methods.

**Table G.5:** Performance Metrics Comparison for Disease-Phenotype Association. Bold values indicate the highest performance among the three methods.

| Test Case | *MedGraphNet* | SRG | Multimodal Fusion |
| --- | --- | --- | --- |
| Disease Isolated Node | **0.90** | 0.33 | 0.89 |
| Phenotype Isolated Node | **0.94** | 0.89 | 0.78 |
| Random Split | 0.90 | **0.96** | 0.93 |

**Table G.6:** Ablation Study: AUC Comparison for All Associations. Bold values indicate the highest performance among the three methods.

| Association | *MedGraphNet* | *MedGraphNet*-Random |
| --- | --- | --- |
| Gene-Drug | **0.91** | 0.88 |
| Disease-Gene | **0.94** | 0.92 |
| Disease-Drug | **0.94** | 0.91 |
| Phenotype-Gene | **0.84** | 0.80 |
| Gene-Drug | **0.91** | 0.88 |
| Disease-Phenotype | **0.90** | 0.86 |

**Figure G.1:**
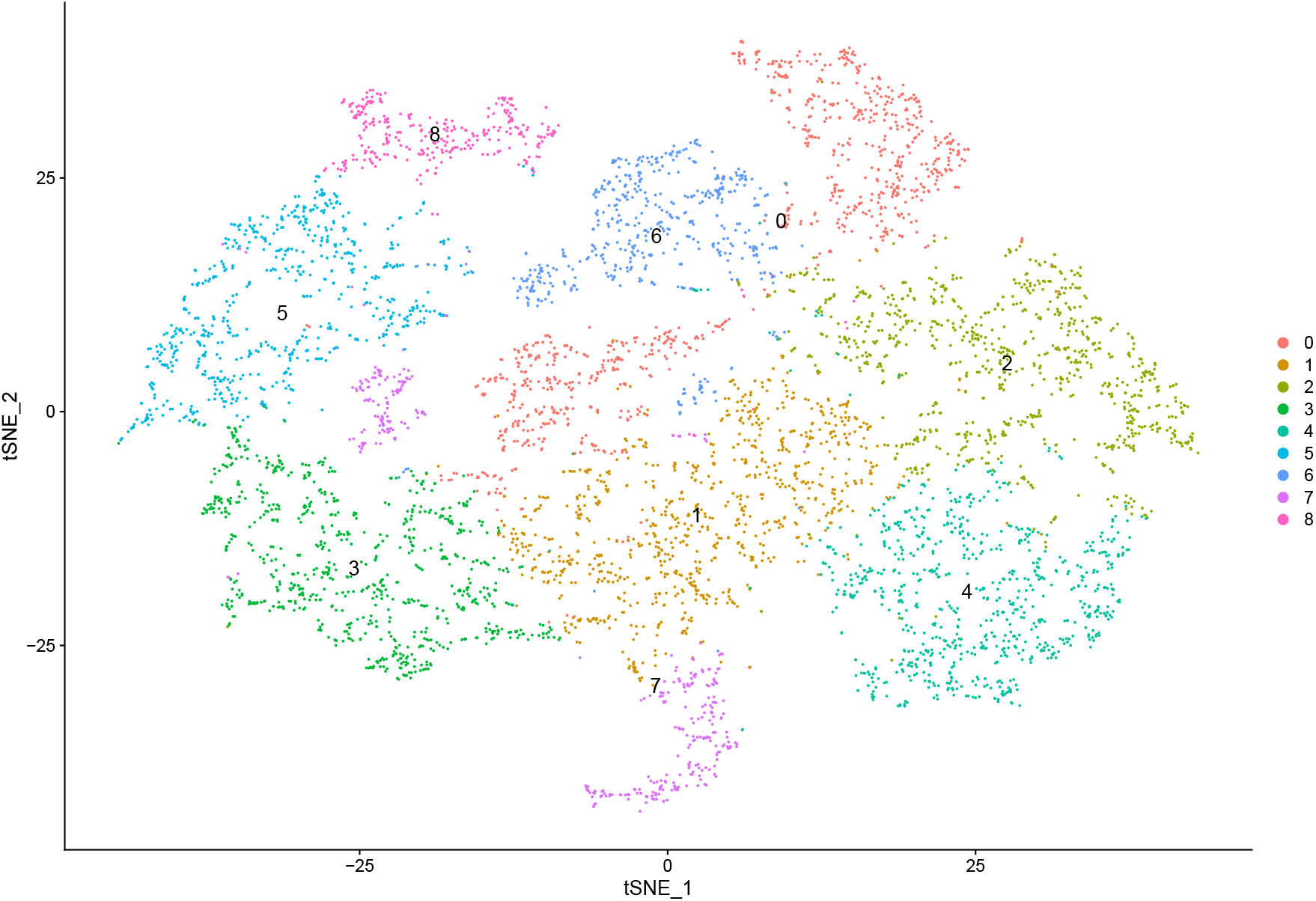
t-SNE Plot of Disease Node Embeddings.

**Figure G.2:**
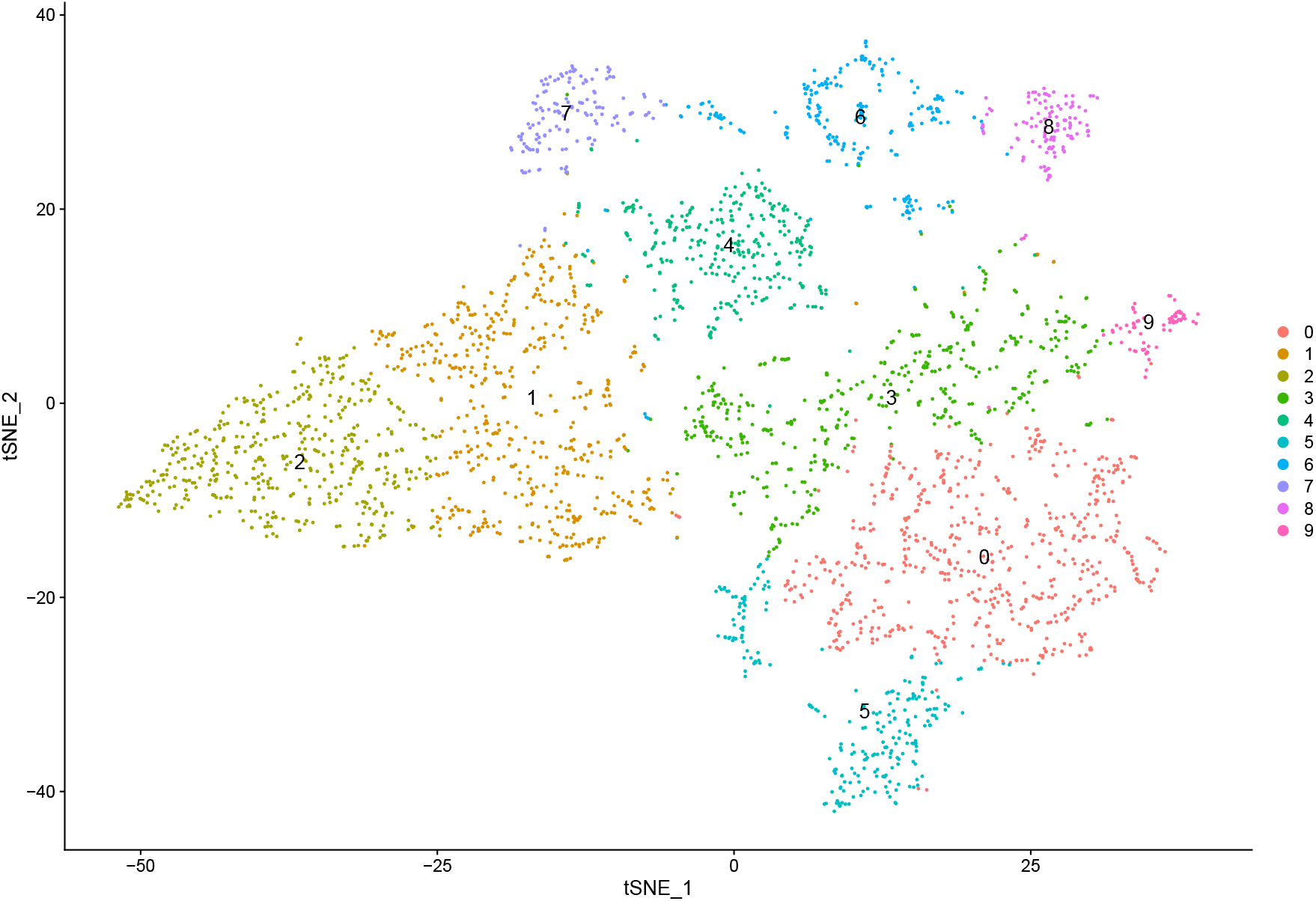
t-SNE Plot of Drug Node Embeddings.

**Figure G.3:**
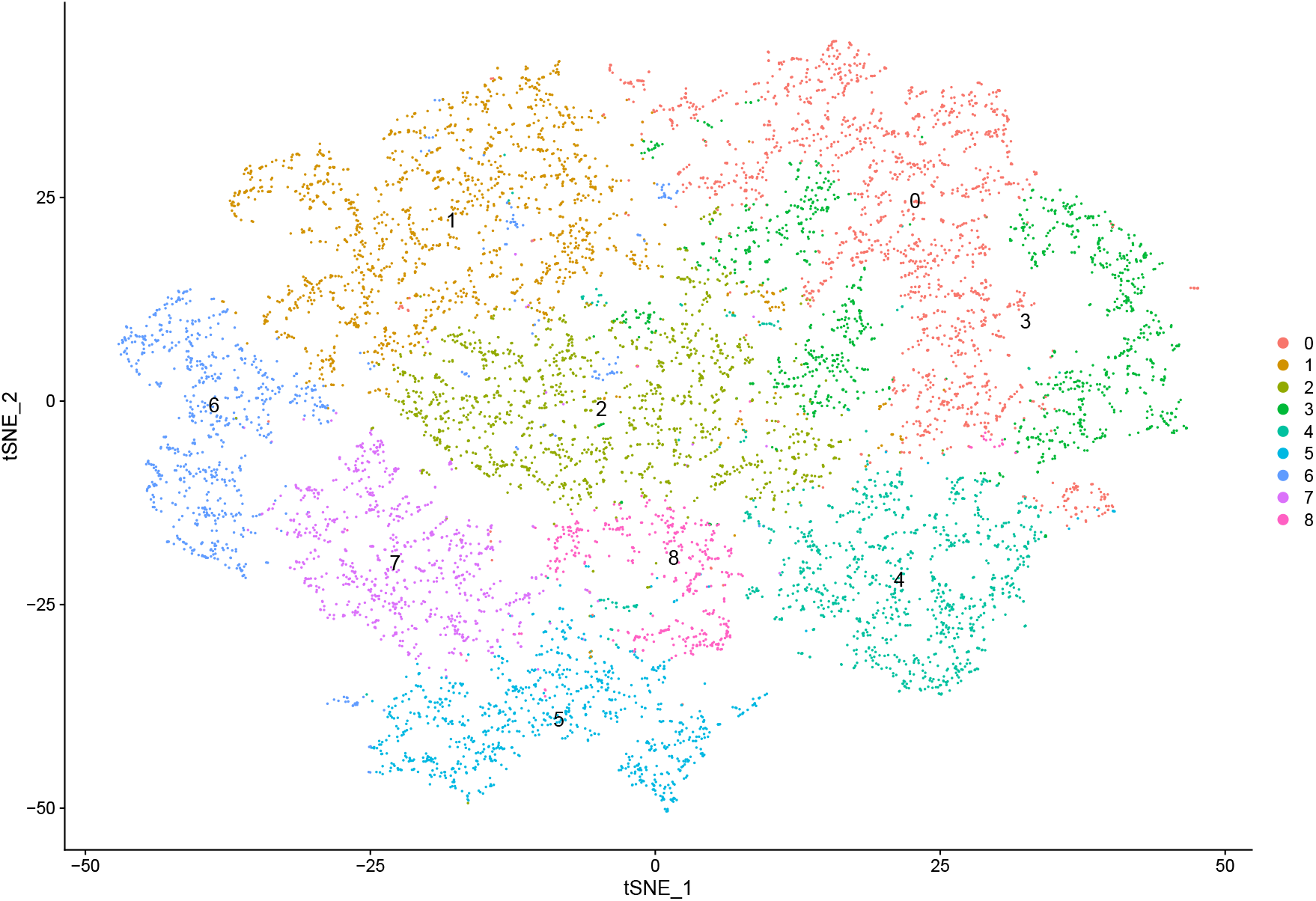
t-SNE Plot of Phenotype Node Embeddings.

## Footnotes

1 https://www.mdpi.com/1099-4300/25/6/909

## References

[1] Davide Angioni, Jeremy Raffin, Pierre-Jean Ousset, Julien Delrieu, and Philipe de Souto Barreto. Fatigue in alzheimer’s disease: Biological basis and clinical management—a narrative review. Aging Clinical and Experimental Research, 35(10):1981– 1989, 2023.

[2] Masaki Asada, Makoto Miwa, and Yutaka Sasaki. Integrating heterogeneous knowledge graphs into drug–drug interaction extraction from the literature. Bioinformatics, 39 (1):btac754, 2023.

[3] Pietro Cinaglia and Mario Cannataro. Identifying candidate gene–disease associations via graph neural networks. Entropy, 25(66):909, June 2023. ISSN 1099-4300. doi: 10.3390/e25060909.

[4] UniProt Consortium. Uniprot: a worldwide hub of protein knowledge. Nucleic acids research, 47(D1):D506–D515, 2019.

[5] Adnan Kivanc Corut and Jason G Wallace. kgwasflow: a modular, flexible, and reproducible snakemake workflow for k-mers-based gwas. G3: Genes, Genomes, Genetics, 14(1):jkad246, 2024.

[6] Pranab Das and Dilwar Hussain Mazumder. An extensive survey on the use of supervised machine learning techniques in the past two decades for prediction of drug side effects. Artificial Intelligence Review, 56(9):9809–9836, 2023.

[7] Sharon L Freshour, Susanna Kiwala, Kelsy C Cotto, Adam C Coffman, Joshua F McMichael, Jonathan J Song, Malachi Griffith, Obi L Griffith, and Alex H Wagner. Integration of the drug–gene interaction database (dgidb 4.0) with open crowdsource efforts. Nucleic acids research, 49(D1):D1144–D1151, 2021.

[8] Alba Gutíerrez-Sacristán, Solene Grosdidier, Olga Valverde, Marta Torrens, A’lex Bravo, Janet Pinero, Ferran Sanz, and Laura I Furlong. Psygenet: a knowledge platform on psychiatric disorders and their genes. Bioinformatics, 31(18):3075–3077, 2015.

[9] Denise Harold, Richard Abraham, Paul Hollingworth, Rebecca Sims, Amy Gerrish, Marian L Hamshere, Jaspreet Singh Pahwa, Valentina Moskvina, Kimberley Dowzell, Amy Williams, et al. Genome-wide association study identifies variants at clu and picalm associated with alzheimer’s disease. Nature genetics, 41(10):1088–1093, 2009.

[10] Barton R Harris, Mark A Prendergast, D Alex Gibson, D Trent Rogers, John A Blanchard, Robert C Holley, May C Fu, Stewart R Hart, Norman W Pedigo, and John M Littleton. Acamprosate inhibits the binding and neurotoxic effects of trans-acpd, suggesting a novel site of action at metabotropic glutamate receptors. Alcoholism: Clinical and Experimental Research, 26(12):1779–1793, 2002.

[11] Paul Hollingworth, Denise Harold, Rebecca Sims, Amy Gerrish, Jean-Charles Lambert, Minerva M Carrasquillo, Richard Abraham, Marian L Hamshere, Jaspreet Singh Pahwa, Valentina Moskvina, et al. Common variants at abca7, ms4a6a/ms4a4e, epha1, cd33 and cd2ap are associated with alzheimer’s disease. Nature genetics, 43(5):429–435, 2011.

[12] Lawrence S Honig, Bruno Vellas, Michael Woodward, Mercé Boada, Roger Bullock, Michael Borrie, Klaus Hager, Niels Andreasen, Elio Scarpini, Hong Liu-Seifert, et al. Trial of solanezumab for mild dementia due to alzheimer’s disease. New England Journal of Medicine, 378(4):321–330, 2018.

[13] Maria Jackson, Leah Marks, Gerhard HW May, and Joanna B Wilson. The genetic basis of disease. Essays in biochemistry, 62(5):643–723, 2018.

[14] Ala Jararweh, Oladimeji Macaulay, David Arredondo, Olufunmilola Oyebamiji, Luis E Tafoya, Kushal Virupakshappa, and Avinash Sahu. Unveiling zero shot prediction for gene attributes through interpretable ai. In ICLR 2024 Workshop on Machine Learning for Genomics Explorations.

[15] Thorlakur Jonsson, Hreinn Stefansson, Stacy Steinberg, Ingileif Jonsdottir, Palmi V Jonsson, Jon Snaedal, Sigurbjorn Bjornsson, Johanna Huttenlocher, Allan I Levey, James J Lah, et al. Variant of trem2 associated with the risk of alzheimer’s disease. New England Journal of Medicine, 368(2):107–116, 2013.

[16] Sunghwan Kim, Jie Chen, Tiejun Cheng, Asta Gindulyte, Jia He, Siqian He, Qingliang Li, Benjamin A Shoemaker, Paul A Thiessen, Bo Yu, et al. Pubchem 2019 update: improved access to chemical data. Nucleic acids research, 47(D1):D1102–D1109, 2019.

[17] David S Knopman, Rosebud O Roberts, Yonas E Geda, Bradley F Boeve, V Shane Pankratz, Ruth H Cha, Eric G Tangalos, Robert J Ivnik, and Ronald C Petersen. Association of prior stroke with cognitive function and cognitive impairment: a population-based study. Archives of Neurology, 66(5):614–619, 2009.

[18] Sebastian Köhler, Michael Gargano, Nicolas Matentzoglu, Leigh C Carmody, David Lewis-Smith, Nicole A Vasilevsky, Daniel Danis, Ganna Balagura, Gareth Baynam, Amy M Brower, et al. The human phenotype ontology in 2021. Nucleic acids research, 49(D1):D1207–D1217, 2021.

[19] Michael Kuhn, Monica Campillos, Ivica Letunic, Lars Juhl Jensen, and Peer Bork. A side effect resource to capture phenotypic effects of drugs. Molecular systems biology, 6(1):343, 2010.

[20] MeiYee Law and David R Shaw. Mouse genome informatics (mgi) is the international resource for information on the laboratory mouse. Eukaryotic Genomic Databases: Methods and Protocols, pages 141–161, 2018.

[21] Jure Leskovec and Andrej Krevl. SNAP Datasets: Stanford large network dataset collection. http://snap.stanford.edu/data, June 2014.

[22] Wenjun Li, Wanjun Ma, Mengyun Yang, and Xiwei Tang. Drug repurposing based on the dtd-gnn graph neural network: revealing the relationships among drugs, targets and diseases. BMC genomics, 25, 2024.

[23] Yuxing Liao, Jing Wang, Eric J Jaehnig, Zhiao Shi, and Bing Zhang. Webgestalt 2019: gene set analysis toolkit with revamped uis and apis. Nucleic acids research, 47(W1): W199–W205, 2019.

[24] Yahui Long, Yu Zhang, Min Wu, Shaoliang Peng, Chee Keong Kwoh, Jiawei Luo, and Xiaoli Li. Heterogeneous graph attention networks for drug virus association prediction. Methods, 198:11–18, 2022.

[25] Qingsong Lv, Ming Ding, Qiang Liu, Yuxiang Chen, Wenzheng Feng, Siming He, Chang Zhou, Jianguo Jiang, Yuxiao Dong, and Jie Tang. Are we really making much progress? revisiting, benchmarking and refining heterogeneous graph neural networks. In Proceedings of the 27th ACM SIGKDD conference on knowledge discovery & data mining, pages 1150–1160, 2021.

[26] Natalya C Maisel, Janet C Blodgett, Paula L Wilbourne, Keith Humphreys, and John W Finney. Meta-analysis of naltrexone and acamprosate for treating alcohol use disorders: when are these medications most helpful? Addiction, 108(2):275–293, 2013.

[27] Janet Pinero, Alex Bravo, Nuŕia Queralt-Rosinach, Alba Gutiérrez-Sacristán, Jordi Deu-Pons, Emilio Centeno, Javier García-García, Ferran Sanz, and Laura I Furlong. Disgenet: a comprehensive platform integrating information on human disease-associated genes and variants. Nucleic acids research, page gkw943, 2016.

[28] Janet Pinero, Juan Manuel Ramírez-Anguita, Josep Saüch-Pitarch, Francesco Ronzano, Emilio Centeno, Ferran Sanz, and Laura I Furlong. The disgenet knowledge platform for disease genomics: 2019 update. Nucleic acids research, 48(D1):D845– D855, 2020.

[29] Ahmet Sureyya Rifaioglu, Heval Atas, Maria Jesus Martin, Rengul Cetin-Atalay, Volkan Atalay, and Tunca Dŏgan. Recent applications of deep learning and machine intelligence on in silico drug discovery: methods, tools and databases. Briefings in bioinformatics, 20(5):1878–1912, 2019.

[30] Stephen Salloway, S Ferris, A Kluger, R Goldman, T Griesing, D Kumar, and S Richardson. Efficacy of donepezil in mild cognitive impairment: a randomized placebo-controlled trial. Neurology, 63(4):651–657, 2004.

[31] Michael Schlichtkrull, Thomas N Kipf, Peter Bloem, Rianne Van Den Berg, Ivan Titov, and Max Welling. Modeling relational data with graph convolutional networks. In The semantic web: 15th international conference, ESWC 2018, Heraklion, Crete, Greece, June 3–7, 2018, proceedings 15, pages 593–607. Springer, 2018.

[32] Lynn M Schriml, James B Munro, Mike Schor, Dustin Olley, Carrie McCracken, Victor Felix, J Allen Baron, Rebecca Jackson, Susan M Bello, Cynthia Bearer, et al. The human disease ontology 2022 update. Nucleic acids research, 50(D1):D1255–D1261, 2022.

[33] Matthias Schwab and Elke Schaeffeler. Pharmacogenomics: a key component of personalized therapy, 2012.

[34] Rogeria Cristina Rangel da Silva, Raquel Luíza Santos de Carvalho, and Marcia Cristina Nascimento Dourado. Deficits in emotion processing in alzheimer’s disease: a systematic review. Dementia & Neuropsychologia, 15:314–330, 2021.

35. Pankhuri Singhal, Shefali Setia Verma, and Marylyn D Ritchie. Gene interactions in human disease studies—evidence is mounting. Annual Review of Biomedical Data Science, 6:377–395, 2023.

[36] Carol Smith, Tariq Rahman, Nicole Toohey, Joseph Mazurkiewicz, Katharine Herrick-Davis, and Milt Teitler. Risperidone irreversibly binds to and inactivates the h5-ht7 serotonin receptor. Molecular pharmacology, 70(4):1264–1270, 2006.

[37] Damian Szklarczyk, Annika L Gable, Katerina C Nastou, David Lyon, Rebecca Kirsch, Sampo Pyysalo, Nadezhda T Doncheva, Marc Legeay, Tao Fang, Peer Bork, et al. The string database in 2021: customizable protein–protein networks, and functional characterization of user-uploaded gene/measurement sets. Nucleic acids research, 49 (D1):D605–D612, 2021.

[38] Vivian Tam, Nikunj Patel, Michelle Turcotte, Yohan Bossé, Guillaume Paré, and David Meyre. Benefits and limitations of genome-wide association studies. Nature Reviews Genetics, 20(8):467–484, 2019.

[39] Farhan Tanvir, Khaled Mohammed Saifuddin, Tanvir Hossain, Arunkumar Bagavathi, and Esra Akbas. Hetrinet: Heterogeneous graph triplet attention network for drug-target-disease interaction. arXiv preprint arXiv:2312.00189, 2023.

[40] Pierre N Tariot, Martin R Farlow, George T Grossberg, Stephen M Graham, Scott McDonald, Ivan Gergel, Memantine Study Group, Memantine Study Group, et al. Memantine treatment in patients with moderate to severe alzheimer disease already receiving donepezil: a randomized controlled trial. Jama, 291(3):317–324, 2004.

[41] Xiao Wang, Houye Ji, Chuan Shi, Bai Wang, Yanfang Ye, Peng Cui, and Philip S Yu. Heterogeneous graph attention network. In The world wide web conference, pages 2022–2032, 2019.

[42] David S Wishart, Yannick D Feunang, An C Guo, Elvis J Lo, Ana Marcu, Jason R Grant, Tanvir Sajed, Daniel Johnson, Carin Li, Zinat Sayeeda, et al. Drugbank 5.0: a major update to the drugbank database for 2018. Nucleic acids research, 46(D1): D1074–D1082, 2018.

[43] Mary K Wojczynski and Hemant K Tiwari. Definition of phenotype. Advances in genetics, 60:75–105, 2008.

[44] Richard John Woodman, Bogda Koczwara, and Arduino Aleksander Mangoni. Applying precision medicine principles to the management of multimorbidity: the utility of comorbidity networks, graph machine learning, and knowledge graphs. Frontiers in Medicine, 10:1302844, 2024.

[45] Chuxu Zhang, Dongjin Song, Chao Huang, Ananthram Swami, and Nitesh V Chawla. Heterogeneous graph neural network. In Proceedings of the 25th ACM SIGKDD international conference on knowledge discovery & data mining, pages 793–803, 2019.

[46] Xiao-Meng Zhang, Li Liang, Lin Liu, and Ming-Jing Tang. Graph neural networks and their current applications in bioinformatics. Frontiers in genetics, 12:690049, 2021.

[47] Zehong Zhang, Lifan Chen, Feisheng Zhong, Dingyan Wang, Jiaxin Jiang, Sulin Zhang, Hualiang Jiang, Mingyue Zheng, and Xutong Li. Graph neural network approaches for drug-target interactions. Current Opinion in Structural Biology, 73:102327, 2022.

